## Supplementary Sections for "Towards a decision support tool for intensive care discharge: machine learning algorithm development using electronic health care data from MIMIC-III and Bristol, UK"

### Supplementary online file for: A machine learning approach to intensive care discharge.

Correspondance to:

**C Feature extraction and missing data** **8**

**D Complete case analysis** **10**

**E Implementation in code** **16**

### A Codifying discharge criteria

The original specification of the nurse-led discharge criteria [1] is presented as a check list with some room for user interpretation. For example, one of the check list questions is: *'Urea and electrolytes normal?'* In order to apply the NLD criteria to historical data we needed to codify them and remove any ambiguity. To do so we first split the original criteria into 16 binary tests, where each test places an upper and/or lower bound on a single physiological variable. If the variable value lies within the bounds the test is passed. For the *'Urea and electrolytes'* criterion we defined four separate tests: one for urea and for each of the three electrolytes identified, via clinical consultation, as most relevant to patient discharge (potassium, sodium, creatinine). We took the 'normal' values of these four variables from the bounds specified in the Philips DAR system in use on GICU.

The blood gas test (R2) was originally multi-variable, either testing the arterial blood gases (PaO2, PaCO2) or the *peripheral capillary oxygen saturation* (SpO2) in the absence of an arterial line. To simplify R2 we neglected PaO2 and PaCO2 (which were missing with high frequency), and tested only SpO2 for all patients.

The airway test (R0) checks if the patient has a patent airway. For this test we used the absence of an endotracheal tube (ETT) as a proxy for the airway being patent.

Finally, we neglected the test of urine output since this could not be successfully located in MIMIC (see section B.1).

Following this procedure we obtained the 15 codified binary tests presented in table 1 of the main text, and referred to throughout as the NLD criteria.

We note here that the criterion value of $9g/dL$, above which haemoglobin is deemed acceptable, does not appear to be evidence based. We refer the reader to this discussion [2] on haemoglobin in critical care. More generally this highlights the problematic nature of scoring systems that place fixed boundary constraints on physiological parameters, even when these boundary values are evidence based. We feel that this motivates the power and algorithmic flexibility of a machine learning approach.

### B Data preprocessing

In this section of the supplementary information we cover the details of the data preprocessing that was required prior to analysis. The basics of these procedures are outlined in the main text, and referenced to the relevant part of this section fuller for explanation.

#### B.1 Data harmonisation and concept mapping

Extensive pre-processing was required for both MIMIC and GICU in order to put the data in a workable format. One reason for this is that both datasets are derived from clinical information systems (CIS) that were never designed for secondary research purposes. Both MIMIC and GICU datasets contained problems with inconsistent naming conventions for physiological parameters. We determined that this problem was less severe in the portion of MIMIC-III derived from the *Metavision* data source (as compared to *Physionet*). Therefore, we restricted our study to this subset of the data. Despite this, extensive concept mapping of disparate *interventionIDs* was required in order to obtain physiological records that were coherent and as close to complete as possible. It was not possible to identify a coherent measure of urine output in MIMIC-III. This is an issue that has previously been discussed on the forums accompanying the MIMIC website. Therefore, we were forced to neglect the discharge criteria based on urine output throughout this study. In general the GICU dataset was easier to work with, in no small part thanks to working with the clinicians who are responsible for administering the associated CIS. When working with both datasets particular care was taken to ensure consistency of measurement units, which differ between the two systems.

#### B.2 Inclusion criteria

As described in the main text we selected study subjects from the MIMIC and GICU datasets according to a number of inclusion criteria. The main goal was to obtain approximately equal sized samples of both patients who were ready for discharge, and those who were not ready for discharge. The GICU data is derived from the clinical information system in use on the general intensive care unit at the Bristol Royal Infirmary. GICU is a combined critical care and high-dependency unit that treats both level 2 and level 3 patients. The MIMIC data is derived from patients admitted to critical care units at the Beth Israel Deaconess Medical Center in Boston, Massachusetts between 2001 and 2012. Initially we applied the following criteria when selecting subjects from the two historical datasets:

- From MIMIC we selected patients from MIMIC-III who were admitted to either medical or surgical intensive care, since this approximates the patient type admitted to general intensive care in Bristol.
- From MIMIC we selected only patient from *Metavision* (see section B.1).
- We only included the first intensive care stay of any given hospital admission. This decision allowed us to calculate readmission rates to ICU. (Readmission was one negative outcome considered following callout.)
- From GICU we included only patients who could be linked to Ward-Watcher (administrative system) via their lifetime ID number.

- From both data sources we selected only patients with a recorded callout (ready for discharge time).
- Furthermore, we discarded any patients (GICU and MIMIC) that had three of more feature variables with no recorded value.
- The two resulting cohorts of study subjects are summarised in table 2 of the main text.

The first step in processing the data was to omit those intensive care stays with no recorded callout. This omission removed any patients who died on ICU, as well as those who should have had a callout recorded but for some reason it was missing in the data. The second step was to define positive and negative outcomes following callout (see main text). Patients with a positive outcome were determined to be ready for discharge at the time of callout. Patients with a negative outcome were determined to be not ready for discharge, either because they were erroneously discharged or because they were discharged in the knowledge that they were not fit (e.g. specialist unit transfer or palliative). For MIMIC and GICU the rate of negative outcomes was 12.57% and 5.83% percent respectively (see the bars at $X_i = 0$ in figure 1).

The next step was to increase the number of instances of NRFD in order to balance to class sizes for training and testing purposes. To produce more instances of NRFD we sampled from earlier in patient stays at times when they were assumed to be not clinically fit for discharge. Heuristically we made the decision that we should sample at least three days prior to callout to ensure no ambiguity in RFD status (one or two days prior to callout it may be the case that patients were RFD but were being held on to conservatively). We also chose to exclude patients who were in the first 24 hours of their ICU stay from this sampling. From figure 1 it is clear that, given the distribution in the length of patient stays, there is only so far back in time we can sample and still obtain

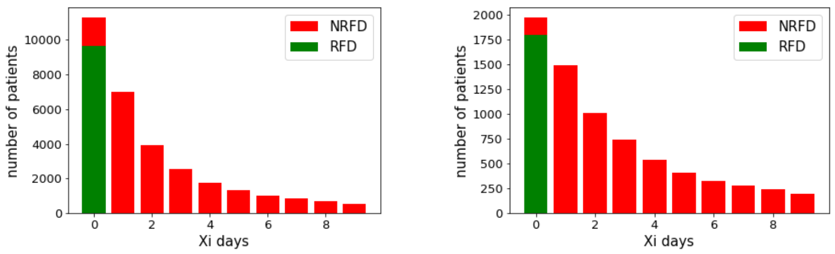

*MIMIC* *GICU*

Figure 1: *Illustrating instances sampled at in integer multiples ($X_i$) of 24 hours prior to callout. Patient callout is at $X_i = 0$. At this time patients with positive outcome are considered ready for discharge (RFD), and patients with negative outcome are considered not ready for discharge (NRFD). At $X_i > 0$ all patients are considered NRFD regardless of subsequent outcome status.*

enough instances of NRFD to balance the class sizes. Therefore, we sampled at time points from to three to eight days prior to callout ($X_i = [3, 4, 5, 6, 7, 8]$). The reason for sampling in exact multiples of 24 hours prior to callout was to ensure the removal of time of day effects from the data (see section B.3). An alternative strategy would be to sample and classify patients at a set time point each day. This strategy would have the benefit of preserving the time relationship of certain daily activities. For example, daily bloods are likely done in the morning. Our strategy, of sampling in 24 hour multiples prior to callout may introduce more variability in the time since last bloods, and possibly other measurements/interventions. However, given that we wanted to work towards a classifier that is not constrained by activities (such as discharge decisions) occurring at fixed times of day, we feel that the chosen sampling strategy is justified.

Sampling patient stays at $X_i = [3, 4, 5, 6, 7, 8]$ prior to callout yielded 8425 and 3174 instances for MIMIC and GICU respectively. We then combined these extra NRFD instances with the original instances (RFD and NRFD) sampled at time of callout ($X_i = 0$), and re-sampled the larger class to obtain a 50:50

ratio of RFD:NRFD. This re-sampling was conducted with the Python method: *SFrame.sample(fraction,seed).* The resulting datasets for MIMIC and GICU contained 7592 and 1870 instances respectively. The subsequent removal of instances with missing data (see section C) reduced these to 5038 and 1851.

##### B.3 Time of day effects

Discharge decisions are invariability made in the morning (figure 2) after discussion at the morning hand over meeting and subsequent ward round. There are non-random temporal signals in the value of certain physiological variables (figure 3). Therefore, it is possible to train a reasonable classifier to identify readiness for discharge by recognising non-random variation in variable values due to time of day. For example: blood pressure is lower than average $\rightarrow$ it must be morning time $\rightarrow$ this patient is more likely to be RFD. This highlights a problem in identifying when patients are actually physically fit for discharge, versus when they are declared so. It also means we must be careful when introducing extra instances of the negative class by sampling from earlier in patient stays - this is why we always sampled in integer multiples of 24 hours prior to callout.

The frequency with which the callout decision is taken around 11am (figure 2) highlights one important way that a decision support system could improve upon ICU discharge practice: by prompting clinicians to consider discharge at other, non-specified time points during the day.

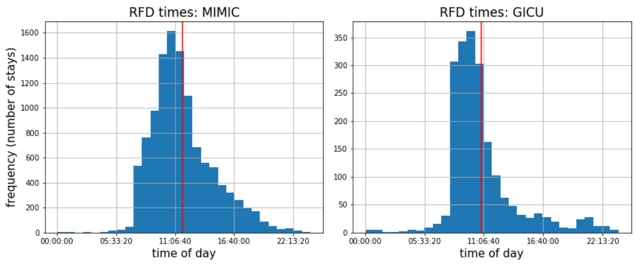

Figure 2: *Histogram of the time of day at which patients were declared ready for discharge (callout time) for: MIMIC (left) and GICU (right). The red line indicates the mean of the distribution.*

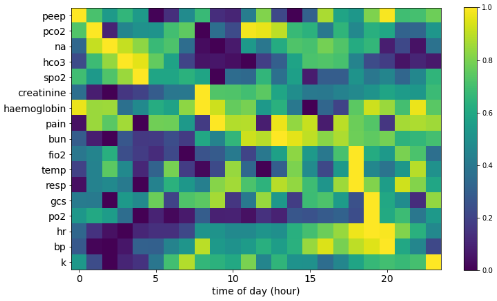

Figure 3: *Variation in average physiological variable values by time of day for the GICU dataset. We calculated the average value for a number of physiological variables at each hour of the day over all patients in the GICU dataset. The values for each variable are than scaled between 0 and 1 to be displayed on the heatmap. Certain temporal patterns become visible. For example, heart rate (HR) and blood pressure (BP) tend to be higher in the evening.*

### C Feature extraction and missing data

We constructed feature matrices for each cohort such that the NLD criteria could easily be tested, and machine learning classifiers could be trained on the same data. In these feature matrices each row represents an instance of either RFD of NRFD, and each column represents a feature. Each of the NLD criteria tests the numeric value of a single physiological parameter. Certain criteria test whether the parameter lies within specified lower and upper bounds, while others impose only a single (lower or upper) bound (see table 1). Therefore, to evaluate a criterion with two bounds requires two feature columns: one giving the minimum value of the relevant parameter, and the other giving the maximum. A single bound criterion requires only one feature column. For example, the features 'resp_min' and 'resp_max' are required for test R4, whereas to test R1 the feature 'fio2_max' alone is sufficient.

We calculated the required minima and maxima of the physiological parameters over four-hour sample windows, as specified by the original NLD criteria. For the original instances (RFD and NRFD) this window constituted the four hours immediately preceding callout. For the extra NRFD instances this was the four hour window immediately preceding the earlier sample time ($X_i$ days prior to callout). In both cases the resulting feature columns contained large amounts of missing data where variables had not been recorded during the specified window, as shown in table 1. This problem was worse with those variables derived from laboratory tests (such as 'bun'), which are infrequently collected (approximately once every 24 hours). To replace some of the missing values the sample window was extended from four to 36 hours. If a missing variable was recorded during this extended window, the latest recorded value was taken for both the minimum and maximum. Clinically it was expected that the main source of missing data is that a variable is less likely to be recorded if it is not of clinical concern. In such cases the extended sample window represents a conservative estimation of a patient's physiological state, by taking parameter values at the last time when they were of concern to clinicians. For the five variables corresponding to double-bound NLD criteria, but with large amounts of missing data (pain, k, bun, creatinine, na) the decision was taken to produce only a single feature for each, rather than two, by taking only the final recorded value. This reduction avoided significant value duplication between feature columns.

| Variables | MIMIC | | GICU | |
|---|---|---|---|---|
| | at CALLOUT | Xi days prior to CALLOUT | at CALLOUT | Xi days prior to CALLOUT |
| *hr_max* | 0.002 (0.000) | 0.007 (0.005) | 0.018 (0.003) | 0.195 (0.194) |
| *hr_min* | 0.002 (0.000) | 0.007 (0.005) | 0.018 (0.003) | 0.195 (0.194) |
| *spo2_max* | 0.011 (0.001) | 0.009 (0.005) | 0.027 (0.004) | 0.202 (0.195) |
| *spo2_min* | 0.011 (0.001) | 0.009 (0.005) | 0.027 (0.003) | 0.202 (0.195) |
| *airway* | 0.985 (0.000) | 0.511 (0.000) | 0.057 (0.000) | 0.215 (0.000) |
| *resp_max* | 0.006 (0.001) | 0.009 (0.006) | 0.057 (0.007) | 0.203 (0.195) |
| *resp_min* | 0.006 (0.001) | 0.009 (0.006) | 0.057 (0.007) | 0.203 (0.195) |
| *bp_min* | 0.007 (0.000) | 0.008 (0.005) | 0.069 (0.039) | 0.254 (0.235) |
| *bp_max* | 0.007 (0.000) | 0.008 (0.005) | 0.069 (0.039) | 0.254 (0.235) |
| *temp_min* | 0.077 (0.002) | 0.079 (0.026) | 0.113 (0.004) | 0.229 (0.197) |
| *temp_max* | 0.077 (0.002) | 0.079 (0.026) | 0.113 (0.004) | 0.229 (0.197) |
| *gcs_max* | 0.106 (0.002) | 0.086 (0.011) | 0.0121 (0.005) | 0.233 (0.196) |
| *gcs_min* | 0.106 (0.002) | 0.086 (0.011) | 0.0121 (0.005) | 0.233 (0.196) |
| *pain* | 0.461 (0.140) | 0.466 (0.152) | 0.194 (0.033) | 0.507 (0.372) |
| *hco3* | 0.885 (0.019) | 0.847 (0.037) | 0.490 (0.075) | 0.312 (0.152) |
| *fio2* | 0.932 (0.686) | 0.380 (0.264) | 0.767 (0.402) | 0.455 (0.320) |
| *haemoglobin* | 0.894 (0.024) | 0.878 (0.042) | 0.975 (0.016) | 0.803 (0.026) |
| *k* | 0.875 (0.017) | 0.810 (0.034) | 0.975 (0.014) | 0.805 (0.028) |
| *bun* | 0.884 (0.019) | 0.854 (0.038) | 0.975 (0.013) | 0.803 (0.025) |
| *creatinine* | 0.884 (0.019) | 0.854 (0.037) | 0.975 (0.015) | 0.805 (0.027) |
| *na* | 0.875 (0.017) | 0.822 (0.034) | 0.975 (0.013) | 0.803 (0.025) |

Table 1: *Summary of missing data in the MIMIC and GICU feature matrices. Values are provided for the samples drawn at time of callout, and samples drawn prior to callout (at $X_i$=[3,4,5,6,7,8]). The main values are the fraction of instances with no value recorded during the four hour window. Values in brackets give fraction of missing data when sample window is extended to 36 hours (see text).*

### D Complete case analysis

In medical data there is likely to be meaning in missingness [3] i.e. data is not missing at random but often due to some other reason. For historical data it is not easy to discern the reasons for missing data, but a naive treatment of missingness is liable to bias the dataset and also to lose valuable information from which to do statistical learning. In the main text we presented results produced from an imputed version of the dataset. To impute the values of missing data entries we used k-nearest neighbours imputation (with $k = 5$). To do this a k-nearest neighbours model was constructed from the complete cases (instances with no missing data). This model was then queried and the mean feature value of the $k = 5$ nearest-neighbours of each instance was taken to replace any missing feature values for that instance. In this section we reproduce the results from the main text but using a complete case analysis with all patients with missing data entries removed. In this way we investigate the sensitivity our results to the missing data values and the k-nn imputation strategy. Beyond this, the optimal treatment of missing data represents a significant avenue for development prior to the deployment of our methodology as a real-time decision support tool.

The removal of the entries with missing data (i.e. complete case analysis) does not change the following key results:

- The random forest and logistic classifiers perform better than the original NLD criteria.
- Weighting the NLD criteria by the feature importance of the logistic classifier improves performance.
- Broadly, which features are most predictive of readiness for discharge remains unchanged, and is consistent between classifiers (Spearman's rank correlation coefficient 0.800 here, 0.761 for imputed data set).

The main difference introduced when moving from the complete case analysis to the imputed data set is slight a drop in performance across the board for the MIMIC cohort. On aggregate it becomes harder to identify MIMIC patients that are ready for discharge when those with missing entries are included. For GICU there is a slight change in the opposite direction. The NLD, random forest and logistic classifier all perform slightly better when GICU patients with missing data entries are included. These performance changes suggest non-random structures in the way that data entries are missing, which likely relate to differences in clinical practice. For example it may be that MIMIC patients that are *nearly* ready for discharge a monitored less closely than more sick patients and therefore have more missing data. The inclusion of patients with missing data would introduce more of these ambiguous cases, which are nearly

| | Predicted RFD | Predicted NRFD |
|---|---|---|
| RFD | 11 | 2496 |
| NRFD | 2 | 2529 |

Table 2: *Confusion matrix for NLD criteria applied to MIMIC feature matrix, with missing data entries removed (complete case analysis).*

| | Predicted RFD | Predicted NRFD |
|---|---|---|
| RFD | 60 | 871 |
| NRFD | 5 | 915 |

Table 3: *Confusion matrix for NLD criteria applied to GICU feature matrix, with missing data entries removed (complete case analysis).*

| | Predicted RFD | Predicted NRFD |
|---|---|---|
| RFD | 75 | 6563 |
| NRFD | 8 | 6597 |

Table 4: *Confusion matrix for NLD criteria applied to MIMIC feature matrix, with missing data filled using k-nearest neighbours. Equivalent to table 2.*

| | Predicted RFD | Predicted NRFD |
|---|---|---|
| RFD | 117 | 1644 |
| NRFD | 7 | 1740 |

Table 5: *Confusion matrix for NLD criteria applied to GICU feature matrix, with missing data filled using k-nearest neighbours. Equivalent to table 3.*

RFD but still being held in critical care. Although the performance changes do not alter the conclusions drawn from the results in the main text, they do suggest that strategies for dealing with missing data should be developed in accordance with on-site clinical practice when deploying such a tool in order to achieve optimal classifications.

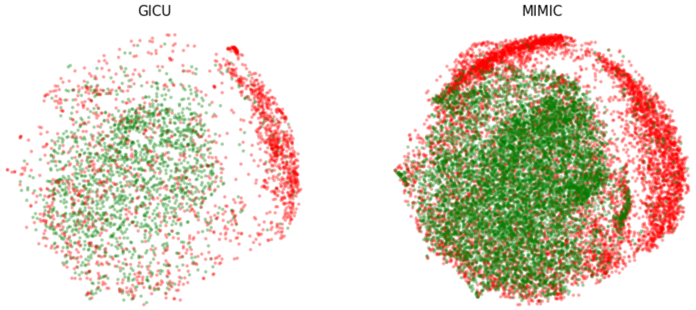

Figure 4: *t-SNE embedding [4] of the feature space for GICU (left), and MIMIC (right), for the imputed dataset. Green and red points indicate instances of RFD and NRFD respectively. The more similar two instances (in terms of feature values), the closer together they appear in the embedding space.*

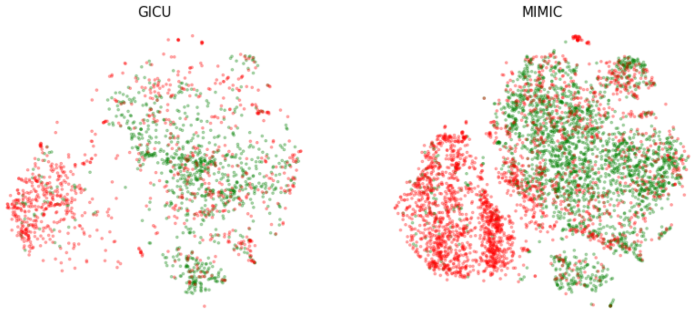

Figure 5: *Equivalent to figure 4, but for the complete case data set.*

| | | NLD | NLD_weighted | LC | RF | LC_extended | RF_extended |
|---|---|---|---|---|---|---|---|
| GICU | AUROC | 0.7802 (0.0173) | 0.8058 (0.0170) | 0.8680 (0.0142) | 0.8544 (0.0148) | **0.8730 (0.0139)** | 0.8619 (0.0146) |
| | Accuracy | 0.7110 (0.0297) | 0.7582 (0.0355) | **0.8284 (0.0501)** | 0.8215 (0.0471) | 0.8279 (0.0485) | 0.8198 (0.0479) |
| | F1 | 0.7366 (0.0176) | 0.7543 (0.0241) | 0.8007 (0.0145) | 0.7966 (0.0166) | **0.8010 (0.0144)** | 0.7954 (0.0184) |
| | Specificity | 0.7000 (0.0000) | 0.7000 (0.0000) | 0.7000 (0.0000) | 0.7000 (0.0000) | 0.7000 (0.0000) | 0.7000 (0.0000) |
| | pAUROC | 0.1412 (0.0096) | 0.1426 (0.0120) | 0.1920 (0.0099) | 0.1815 (0.0103) | **0.1951 (0.0098)** | 0.1853 (0.0106) |
| | Brier | 0.2694 (0.0085) | 0.2419 (0.0124) | 0.1598 (0.0085) | 0.1675 (0.0076) | **0.1578 (0.0085)** | 0.1623 (0.0080) |
| | Sensitivity | 0.7302 (0.0281) | 0.7750 (0.0405) | **0.8750 (0.0232)** | 0.8640 (0.0255) | 0.8732 (0.0218) | 0.8615 (0.0298) |
| MIMIC | AUROC | 0.7544 (0.0097) | 0.8686 (0.0076) | 0.8880 (0.0137) | 0.8898 (0.0143) | 0.8910 (0.0131) | **0.8990 (0.0125)** |
| | Accuracy | 0.6747 (0.0148) | 0.8545 (0.0537) | 0.8641 (0.0607) | 0.8651 (0.0617) | 0.8646 (0.0609) | **0.8775 (0.0654)** |
| | F1 | 0.7196 (0.0099) | 0.8203 (0.0091) | 0.8248 (0.0155) | 0.8257 (0.0170) | 0.8254 (0.0153) | **0.8339 (0.0141)** |
| | Specificity | 0.7000 (0.0000) | 0.7000 (0.0000) | 0.7000 (0.0000) | 0.7000 (0.0000) | 0.7000 (0.0000) | 0.7000 (0.0000) |
| | pAUROC | 0.1315 (0.0052) | 0.1816 (0.0064) | 0.1979 (0.0117) | 0.1987 (0.0122) | 0.2017 (0.0109) | **0.2047 (0.0114)** |
| | Brier | 0.2443 (0.0039) | 0.1551 (0.0068) | 0.1319 (0.0082) | 0.1343 (0.0074) | 0.1303 (0.0078) | **0.1279 (0.0066)** |
| | Sensitivity | 0.6724 (0.0184) | 0.9070 (0.0145) | 0.9221 (0.0224) | 0.9230 (0.0268) | 0.9225 (0.0234) | **0.9405 (0.0217)** |

Table 6: *Performance summary for the complete case analysis, equivalent to table 3 in the main text. All scores are averaged over 100 train-test data splits and given as: mean (standard deviation). All metrics other than AUROC and Brier score are evaluated at a specificity of 0.7, using linear interpolation to estimate this operating point in ROC-space.* $NLD_{weighted}$ *are the NLD criteria, weighted by feature importances from the logistic classifier (table 4).* $LC_{extended}$ *and* $RF_{extended}$ *are the machine learning classifiers with extended feature sets.*

| | FPR | NLD | NLD_weighted | LR | RF | LR_extended | RF_extended |
|---|---|---|---|---|---|---|---|
| GICU | 0.1 | 0.4231 (0.0330) | 0.4259 (0.0341) | 0.6002 (0.0367) | 0.5600 (0.0375) | **0.6274 (0.0327)** | 0.5755 (0.0406) |
| | 0.2 | 0.6109 (0.0234) | 0.6334 (0.0256) | **0.7937 (0.0211)** | 0.7633 (0.0308) | 0.7915 (0.0221) | 0.7744 (0.0283) |
| | 0.3 | 0.7426 (0.0166) | 0.8098 (0.0263) | 0.8870 (0.0171) | 0.8860 (0.0196) | 0.8767 (0.0196) | **0.8909 (0.0185)** |
| | 0.4 | 0.8204 (0.0152) | 0.9244 (0.0125) | 0.9417 (0.0104) | 0.9418 (0.0093) | 0.9404 (0.0123) | **0.9473 (0.0108)** |
| | 0.5 | 0.8884 (0.0114) | 0.9514 (0.0083) | 0.9706 (0.0073) | 0.9653 (0.0072) | **0.9766 (0.0074)** | 0.9711 (0.0068) |
| | 0.6 | 0.9253 (0.0092) | 0.9752 (0.0050) | 0.9860 (0.0047) | 0.9815 (0.0061) | **0.9877 (0.0042)** | 0.9824 (0.0057) |
| | 0.7 | 0.9609 (0.0081) | 0.9804 (0.0047) | 0.9903 (0.0041) | 0.9886 (0.0046) | **0.9918 (0.0037)** | 0.9897 (0.0044) |
| | 0.8 | 0.9791 (0.0047) | 0.9899 (0.0036) | 0.9945 (0.0030) | 0.9929 (0.0035) | **0.9951 (0.0028)** | 0.9937 (0.0034) |
| | 0.9 | 0.9927 (0.0025) | 0.9973 (0.0017) | 0.9957 (0.0025) | 0.9965 (0.0023) | 0.9963 (0.0024) | **0.9970 (0.0022)** |
| | 1 | 1.0000 (0.0000) | 1.0000 (0.0000) | 0.9999 (0.0004) | 1.0000 (0.0000) | 1.0000 (0.0000) | 1.0000 (0.0002) |
| MIMIC | 0.1 | 0.3370 (0.0128) | 0.3740 (0.0185) | 0.4611 (0.0484) | 0.4858 (0.0454) | 0.5449 (0.0482) | **0.5933 (0.0452)** |
| | 0.2 | 0.5296 (0.0110) | 0.6355 (0.0211) | 0.7466 (0.0404) | 0.7465 (0.0442) | 0.7825 (0.0304) | **0.7881 (0.0349)** |
| | 0.3 | 0.6713 (0.0126) | 0.8337 (0.0174) | 0.8827 (0.0282) | 0.8860 (0.0265) | 0.9001 (0.0207) | **0.9049 (0.0210)** |
| | 0.4 | 0.7545 (0.0086) | 0.9295 (0.0091) | 0.9526 (0.0152) | 0.9498 (0.0158) | 0.9590 (0.0111) | **0.9597 (0.0125)** |
| | 0.5 | 0.8309 (0.0096) | 0.9777 (0.0045) | 0.9842 (0.0075) | 0.9853 (0.0065) | 0.9872 (0.0060) | **0.9903 (0.0055)** |
| | 0.6 | 0.8869 (0.0069) | 0.9925 (0.0016) | 0.9943 (0.0035) | 0.9952 (0.0030) | 0.9939 (0.0031) | **0.9973 (0.0026)** |
| | 0.7 | 0.9315 (0.0056) | 0.9950 (0.0013) | 0.9961 (0.0028) | 0.9972 (0.0024) | 0.9961 (0.0024) | **0.9985 (0.0018)** |
| | 0.8 | 0.9659 (0.0036) | 0.9969 (0.0010) | 0.9980 (0.0019) | 0.9985 (0.0018) | 0.9971 (0.0022) | **0.9991 (0.0015)** |
| | 0.9 | 0.9887 (0.0022) | 0.9990 (0.0006) | 0.9993 (0.0011) | 0.9996 (0.0008) | 0.9982 (0.0019) | **0.9997 (0.0007)** |
| | 1 | 1.0000 (0.0000) | 1.0000 (0.0000) | 1.0000 (0.0000) | 1.0000 (0.0000) | 0.9998 (0.0005) | 1.0000 (0.0000) |

Table 7: *Sensitivity values for the various classifiers over a range of false positive rates (FPR), for the* ***imputed data set****. Specificity = 1 - FPR.*

| | FPR | NLD | NLD_weighted | LR | RF | LR_extended | RF_extended |
|---|---|---|---|---|---|---|---|
| GICU | 0.1 | 0.4003 (0.0433) | 0.4117 (0.0564) | 0.5944 (0.0413) | 0.5405 (0.0521) | **0.6142 (0.0431)** | 0.5733 (0.0490) |
| | 0.2 | 0.5928 (0.0389) | 0.6074 (0.0385) | 0.7731 (0.0397) | 0.7470 (0.0414) | **0.7815 (0.0367)** | 0.7492 (0.0400) |
| | 0.3 | 0.7302 (0.0281) | 0.7750 (0.0405) | **0.8750 (0.0232)** | 0.8640 (0.0255) | 0.8732 (0.0218) | 0.8615 (0.0298) |
| | 0.4 | 0.8053 (0.0252) | 0.8899 (0.0241) | 0.9188 (0.0201) | 0.9143 (0.0193) | 0.9222 (0.0199) | **0.9250 (0.0194)** |
| | 0.5 | 0.8735 (0.0213) | 0.9318 (0.0155) | 0.9537 (0.0148) | 0.9478 (0.0146) | **0.9589 (0.0141)** | 0.9589 (0.0148) |
| | 0.6 | 0.9164 (0.0158) | 0.9612 (0.0132) | 0.9734 (0.0110) | 0.9631 (0.0113) | **0.9779 (0.0092)** | 0.9726 (0.0101) |
| | 0.7 | 0.9564 (0.0131) | 0.9744 (0.0086) | 0.9837 (0.0072) | 0.9802 (0.0096) | **0.9868 (0.0066)** | 0.9844 (0.0076) |
| | 0.8 | 0.9764 (0.0077) | 0.9846 (0.0060) | 0.9925 (0.0051) | 0.9885 (0.0064) | **0.9932 (0.0049)** | 0.9907 (0.0049) |
| | 0.9 | 0.9916 (0.0041) | 0.9952 (0.0032) | 0.9973 (0.0026) | 0.9965 (0.0034) | **0.9974 (0.0024)** | 0.9958 (0.0037) |
| | 1 | 1.0000 (0.0000) | 1.0000 (0.0000) | 1.0000 (0.0000) | 1.0000 (0.0000) | 1.0000 (0.0000) | 1.0000 (0.0000) |
| MIMIC | 0.1 | 0.3880 (0.0238) | 0.4841 (0.0392) | 0.6001 (0.0603) | 0.6030 (0.0651) | 0.6211 (0.0598) | **0.6360 (0.0573)** |
| | 0.2 | 0.5473 (0.0208) | 0.7946 (0.0277) | 0.8285 (0.0370) | 0.8217 (0.0402) | 0.8408 (0.0377) | **0.8417 (0.0393)** |
| | 0.3 | 0.6724 (0.0184) | 0.9070 (0.0145) | 0.9221 (0.0224) | 0.9230 (0.0268) | 0.9225 (0.0234) | **0.9405 (0.0217)** |
| | 0.4 | 0.7630 (0.0160) | 0.9635 (0.0085) | 0.9700 (0.0119) | 0.9710 (0.0123) | 0.9710 (0.0118) | **0.9819 (0.0107)** |
| | 0.5 | 0.8433 (0.0117) | 0.9794 (0.0044) | 0.9849 (0.0072) | 0.9884 (0.0058) | 0.9825 (0.0075) | **0.9937 (0.0051)** |
| | 0.6 | 0.8905 (0.0091) | 0.9857 (0.0033) | 0.9905 (0.0054) | 0.9924 (0.0046) | 0.9889 (0.0059) | **0.9962 (0.0036)** |
| | 0.7 | 0.9373 (0.0086) | 0.9901 (0.0032) | 0.9935 (0.0041) | 0.9939 (0.0044) | 0.9914 (0.0051) | **0.9975 (0.0029)** |
| | 0.8 | 0.9655 (0.0053) | 0.9947 (0.0022) | 0.9969 (0.0032) | 0.9975 (0.0030) | 0.9946 (0.0042) | **0.9987 (0.0019)** |
| | 0.9 | 0.9872 (0.0032) | 0.9978 (0.0015) | 0.9987 (0.0019) | 0.9995 (0.0013) | 0.9977 (0.0025) | **0.9997 (0.0010)** |
| | 1 | 1.0000 (0.0000) | 1.0000 (0.0000) | 1.0000 (0.0000) | 1.0000 (0.0000) | 0.9997 (0.0010) | 1.0000 (0.0000) |

Table 8: *Sensitivity values for the various classifiers over a range of false positive rates (FPR), for the* ***complete case data set****. Specificity = 1 - FPR.*

| | Importance (LC) | Importance (RF) | Rank (LC) | Rank (RF) |
|---|---|---|---|---|
| **gcs_min** | 0.1002 (0.0045) | 0.0920 (0.0121) | 0 | 0 |
| **airway** | 0.0874 (0.0105) | 0.0780 (0.0117) | 1 | 1 |
| **bun** | 0.0169 (0.0015) | 0.0106 (0.0018) | 2 | 3 |
| **fio2** | 0.0097 (0.0010) | 0.0152 (0.0019) | 3 | 2 |
| **haemoglobin** | 0.0090 (0.0011) | 0.0093 (0.0018) | 4 | 4 |
| **hr_max** | 0.0045 (0.0020) | 0.0053 (0.0012) | 5 | 5 |
| **resp_max** | 0.0040 (0.0016) | 0.0017 (0.0005) | 6 | 10 |
| **hr_min** | 0.0038 (0.0015) | 0.0031 (0.0008) | 7 | 6 |
| **resp_min** | 0.0025 (0.0015) | 0.0031 (0.0006) | 8 | 7 |
| **spo2_min** | 0.0012 (0.0007) | 0.0005 (0.0002) | 9 | 12 |
| **hco3** | 0.0004 (0.0003) | 0.0002 (0.0001) | 10 | 16 |
| **temp_max** | 0.0002 (0.0006) | 0.0023 (0.0006) | 11 | 8 |
| **na** | 0.0002 (0.0002) | 0.0002 (0.0001) | 12 | 15 |
| **pain** | 0.0001 (0.0001) | 0.0010 (0.0003) | 13 | 11 |
| **temp_min** | 0.0001 (0.0005) | 0.0003 (0.0002) | 14 | 14 |
| **bp_min** | 0.0001 (0.0001) | 0.0005 (0.0002) | 15 | 13 |
| **k** | 0.0000 (0.0000) | 0.0002 (0.0001) | 16 | 17 |
| **creatinine** | 0.0000 (0.0000) | 0.0018 (0.0005) | 17 | 9 |

Table 9: *Feature importances under the complete case analysis, equivalent to table 4 in the main text.*

### E Implementation in code

Data extraction and preprocessing were conducted using SQL. We worked with a local MySQL instance of the MIMIC-III data, and the Bristol (GICU) data was accessed from the backend of the Philips DAR clinical information system via Microsoft SQL Reporting Services on the hospital server. Additional patient demographic information was obtained from the WardWatcher system, which is the ICNARC reporting software in use on the general intensive care unit in Bristol. Entries from WardWatcher were linked to the Philips data via patient lifetime identification numbers. The data were then anonymised by the removal of all sensitive information, including times and date, except for those variables required for the analysis (see main text). Subsequent data processing and analysis was conducted in Python within UH Bristol. We used the machine learning package scikit-learn to train and test classifiers. In particular the method *GridsearchCV* was used to optimise model hyperparamaters via multiple-source cross-validation, optimising for the F1-score. This optimisation was conducted over two folds, where each fold contained only data from a single source (either GICU or MIMIC). The 'best_estimator' parameter of *GridsearchCV* was set to 'True' such that the optimised model was refitted to the full training data (MIMIC and GICU). For the logistic classifier the regularisation hyperparameter 'c' was optimised over a logarithmic range of twenty values between $1 \times 10^{-3}$ and $1 \times 10^{3}$. For the random forest three hyperparameters were optimised over the following ranges: $n\_estimators \in [20, 50, 100]$; $max\_features \in [20, 50, 100]$; $max\_depth \in [4, 5, 6, 7]$. Feature importances were calculated using the *PermutationImportance* method from the package ELI5, whereby the importance is taken as the average loss in performance (AUROC score) when the feature values are randomly permuted.
